# Molecular determinants of RNase MRP specificity and function

**DOI:** 10.1101/2025.01.28.635360

**Authors:** Eric M. Smith, Jimmy Ly, Sofia Haug, Iain M. Cheeseman

## Abstract

RNase MRP and RNase P are evolutionarily related complexes that facilitate rRNA and tRNA biogenesis, respectively. The two enzymes share nearly all protein subunits and have evolutionarily related catalytic RNAs. Notably, RNase P includes a unique subunit, Rpp21, whereas no RNase MRP-specific proteins have been found in humans, limiting molecular analyses of RNase MRP function. Here, we identify the RNase MRP-specific protein, C18orf21/RMRPP1. RMRPP1 and Rpp21 display significant structural homology, but we identify specific regions that drive interactions with their respective complexes. Additionally, we reveal that RNase MRP is required for 40S, but not 60S, ribosome biogenesis uncovering an alternative pathway for ribosome assembly. Finally, we identify Nepro as an essential rRNA processing factor that associates with the RNase MRP complex. Together, our findings elucidate the molecular determinants of RNase MRP function and underscore its critical role in ribosome biogenesis.

## Introduction

Ribosomal RNA (rRNA) is the most abundant cellular RNA molecule with the production and maturation of rRNA being critical for ribosome function^1,2^. In humans, there are four mature rRNAs that are produced from two different non-coding RNA transcripts^3^. The 5S rRNA is transcribed by RNA polymerase III whereas the 5.8S, 28S, and 18S rRNAs are transcribed as a polycistronic transcript by RNA Polymerase I (Fig.1A)^4^. The mature 5.8S, 28S, and 18S rRNAs are formed through endonucleolytic cleavage at specific sites in externally and internally transcribed spacers (ITS) followed by exonucleolytic action^5–7^. The ribonucleoprotein complex RNase MRP acts as a site-specific endonuclease to cleave pre-rRNA at site 2 in the ITS1 (Fig. 1A)^8,9^. The RNase MRP complex is composed of a catalytic RNA subunit, termed RNase MRP RNA, and at least 9 protein subunits (Pop1, Pop5, Rpp38, Rpp30, Rpp29, Rpp25, Rpp20, Rpp14, and Rpp40)^10–18^. The RNase MRP RNA is transcribed from the *RMRP* gene, which is conserved across eukaryotes ^13,19,20^. The importance of RNase MRP is highlighted by the fact that mutations in RNase MRP RNA are embryonic lethal in mouse and result in a variety of human diseases including cartilage hair hypoplasia, metaphyseal dysplasia without hypotrichosis, anauxetic dysplasia, kyphomelic dysplasia, and Omenn syndrome^21–24^.

**Figure 1.**
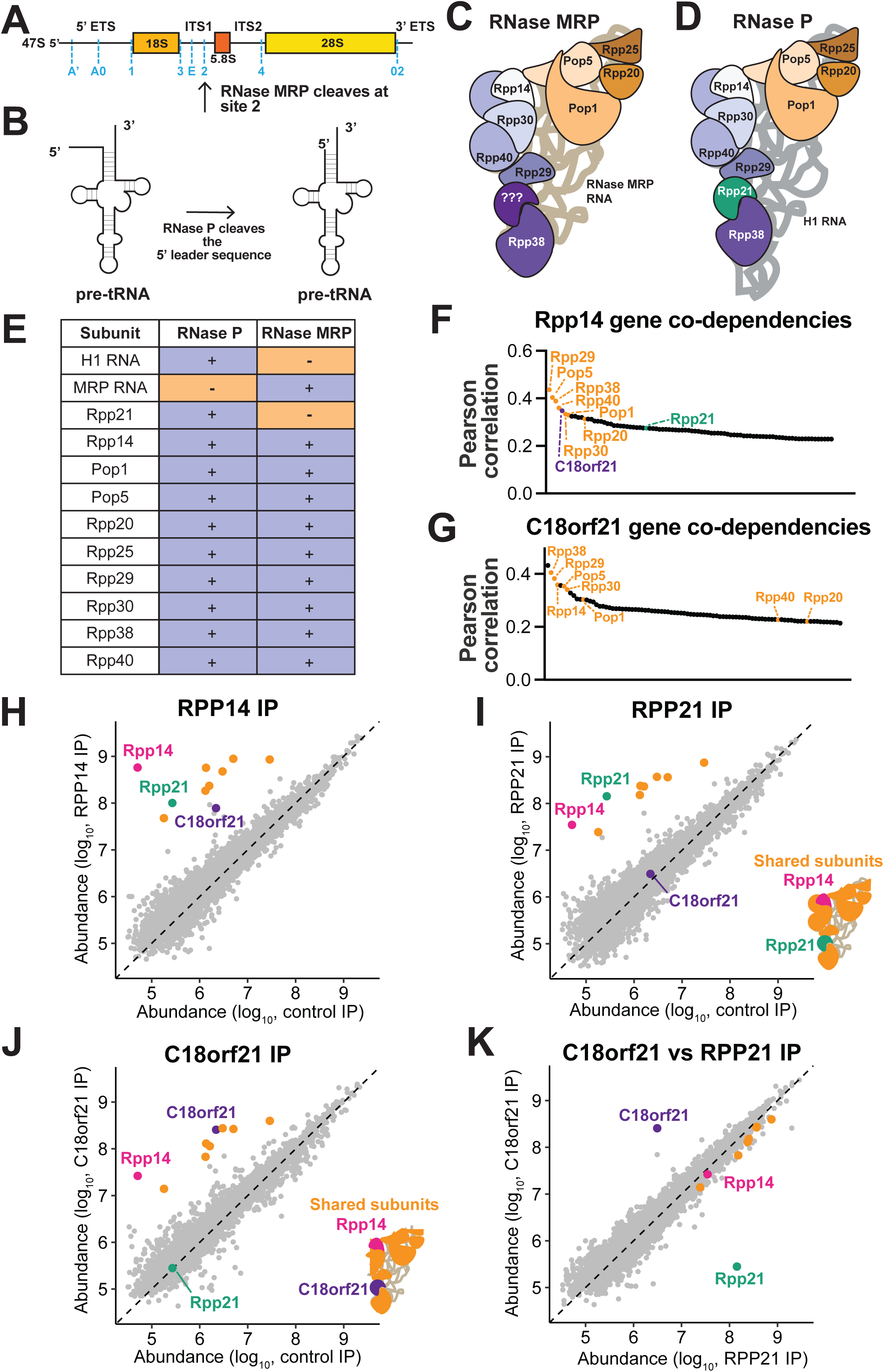
C18orf21 is associated with the shared subunits of the RNase P and RNase MRP complexes. **(A)** A schematic of the 47S rRNA transcript. Endonucleolytic cleavage sites in the externally and internally transcribed spaces (ETS and ITS) are marked in blue. RNase MRP cleaves at site 2 in ITS1. **(B)** A schematic of a precursor tRNA before and after the cleavage of the 5’ leader sequence by RNase P. **(C)** A cartoon representation of RNase MRP. **(D)** A cartoon representation of RNase P. **(E)** A table showing which subunits are found in either the RNase P complex, the RNase MRP complex, or in both complexes. **(F)** A waterfall plot showing the genes with the highest Pearson correlation coefficients to Rpp14. These coefficients are acquired from the CRISPR-Cas9 targeting effect scores in the Depmap database. The shared RNase P and MRP components are shown in orange, C18orf21 is shown in purple, and Rpp21 is shown in green. **(G)** A waterfall plot showing the genes with the highest Pearson correlation of CRISPR-Cas9 based targeting effect scores to C18orf21 in the Depmap database. The shared RNase P and MRP components are shown in orange. **(H-K)** Scatter plots showing the abundance of the proteins detected in the indicated immunoprecipitation-mass spectrometry experiments. The shared components of the RNase P/MRP complexes are highlighted in orange, except Rpp14 which is shown in pink. C18orf21 is highlighted in purple and Rpp21 is highlighted in green.

In addition to acting as an essential enzyme for rRNA processing, RNase MRP is a fascinating example of evolution repurposing a core cellular machine to carry out a novel function. RNase MRP is evolutionarily-related to the ribonucleoprotein complex RNase P ^20,25–27^. Like RNase MRP, RNase P acts as an endonuclease, but cleaves the 5’ leader sequence of pre-tRNAs (Fig. 1B)^28^. Due to its essential function in tRNA biogenesis, RNase P is conserved across all three domains of life^29^. RNase MRP is thought to have evolved in eukaryotes as a result of the duplication of the RNase P RNA gene *RPPH1* (H1 RNA), which gave rise to the *RMRP* gene that encodes the RNase MRP RNA^10,25^. Structural and biochemical studies of *S. cerevisiae* RNase P and RNase MRP have revealed that the catalytic domains and RNA substrate cleavage mechanism are shared between the two RNA subunits, despite differences between the sequences of each RNA^11,12,25,30^. In addition to their similar RNA subunits, RNase P and RNase MRP share common protein subunits (Fig. 1C-E)^10^. In humans, only one protein, Rpp21, has been identified as an RNase P-specific protein^31^. Despite the presence of two RNase MRP-specific subunits in *S. cerevisiae*, no known homologs to these proteins exist in vertebrates and no other RNase MRP-specific proteins have been identified^13,32,33^. Because of the lack of knowledge of a unique protein subunit for the enzyme, the molecular requirements for the activities of RNase MRP in human cells have remained unclear.

Here, we identify the uncharacterized protein C18orf21/RMRPP1 as an RNase MRP-specific protein subunit. This identification allowed us to distinguish RNase MRP from RNase P function at the protein level, define sequence features that confer specificity of C18orf21 and Rpp21 for RNase MRP and RNase P, and reveal a critical role for C18orf21/RMRPP1 in rRNA processing, ribosome biogenesis, and cellular fitness. Finally, we leverage the identification of RMRPP1 to define Nepro as a critical rRNA processing factor through its interaction with the RNase MRP complex. Together, this work reveals key molecular determinants of RNase MRP specificity and function.

## Results

### C18orf21 associates with the shared subunits of the RNase P/MRP complexes

To identify protein subunits that distinguish the functional properties and activities of the RNase MRP complex from the RNase P complex, we evaluated genetic co-dependencies. The use of functional genetics in human cells has enabled effective strategies to identify co-functional genes, including factors that function as part of the same protein complex^34,35^. To identify previously uncharacterized proteins, we conducted an analysis of gene co-essentiality using genome-wide CRISPR-Cas9 based viability screens performed on >1000 different cells lines as part of the Project Achilles Cancer Dependency map^36^. Based on similar patterns of gene requirements across these diverse cell lines, we found that the shared component of the RNase P and RNase MRP complexes, Rpp14, was genetically correlated with the other shared components of the two complexes. Surprisingly, we also found that Rpp14 had a strong co-dependency with the uncharacterized gene C18orf21 (Fig.1F). Reciprocally, we found that C18orf21 displayed similar genetic dependency requirements to 8 out of 9 genes that are shared between the RNase P and RNase MRP complexes (Fig.1G). C18orf21 also displays strongly correlated functional requirements with several genes that are involved in rRNA production (Extended Data Fig.1A). In contrast to the strong correlation of C18orf21 with the components of the RNase P/MRP complexes, the RNase P-specific component, Rpp21, displayed genetic correlations with only two of the shared RNase P/MRP components as well as correlations with several tRNA processing factors (Extended Data Fig.1B). Similarly, recent genome-wide perturbation studies combined with single cell sequencing (Perturb-Seq) identified phenotypic similarities between C18orf21 and genes associated with rRNA biogenesis, based on transcriptome similarities in depleted cells, whereas Rpp21 displayed phenotypic similarities with tRNA processing genes^37^.

Based on the co-functional relationships that we observed between C18orf21 and members of the RNase MRP and RNase P complexes, we next sought to determine if C18orf21 interacts with any of the shared RNase P/MRP components in cells. To identify binding partners for C18orf21, we conducted immunoprecipitation-mass spectrometry experiments (IP-MS) from HeLa cells. In particular, we isolated C-terminally tagged GFP fusion proteins for C18orf21, Rpp14, which acts as a shared subunit of both RNase P and RNase MRP, and the RNase P-specific protein Rpp21. Rpp14 immunoprecipitations isolated each of the nine protein subunits that are shared by RNase P and RNase MRP, as well as isolating both Rpp21 and C18orf21 (Fig.1H). In contrast, we found that Rpp21 purifications isolated all nine of the expected binding partners from the RNase P complex, but did not co-immunoprecipitate with C18orf21 (Fig.1I). Finally, we found that C18orf21 isolated the nine subunits common to both RNase P and RNase MRP, but did not isolate Rpp21 (Fig.1J). Thus, C18orf21 and Rpp21 display mutually exclusive protein interactions, but each protein associates with the shared subunits of the RNase P and RNase MRP complexes (Fig.1K and Extended Data Fig.1C).

### C18orf21 (RMRPP1) acts as an RNase MRP complex-specific protein in cells

Our genetic interaction and IP-MS analyses suggest that C18orf21 and Rpp21 display mutually exclusive interactions with RNase P/MRP. We hypothesized that, as Rpp21 is specific to RNase P, C18orf21 may be specific to RNase MRP such that both of these proteins interact with shared components but not with each other. To evaluate this model further, we next assessed the cellular properties of C18orf21 and Rpp21. Prior work suggested that RNase MRP and RNase P localize to distinct compartments within the nucleus^31,38^. Therefore, we analyzed the cellular localization of C18orf21, the shared RNase P/MRP subunit Rpp14, and the RNase P-specific protein Rpp21 as GFP fusions in HeLa cells. We found that Rpp14 displayed dual localization to the nucleus, nuclear puncta, and to the nucleolus, consistent with the combined reported localization for both RNase P and RNase MRP (Fig.2A). In contrast, Rpp21-GFP localized to nuclear foci, but not the nucleolus, consistent with a specificity for its localization as an RNase P-specific protein (Fig.2A). We found that C18orf21 localized only to the nucleolus, but not throughout the nucleus, consistent with a role in promoting the specific activities of RNase MRP (Fig.2A). To evaluate if C18orf21 is required for the nucleolar localization of other RNase MRP components, we monitored Rpp14 localization in C18orf21 knockout cells. Strikingly, we found that loss of C18orf21 led to a loss of the nucleolar population of Rpp14, but did not disrupt its localization to nuclear foci (Fig.2B).

**Figure 2.**
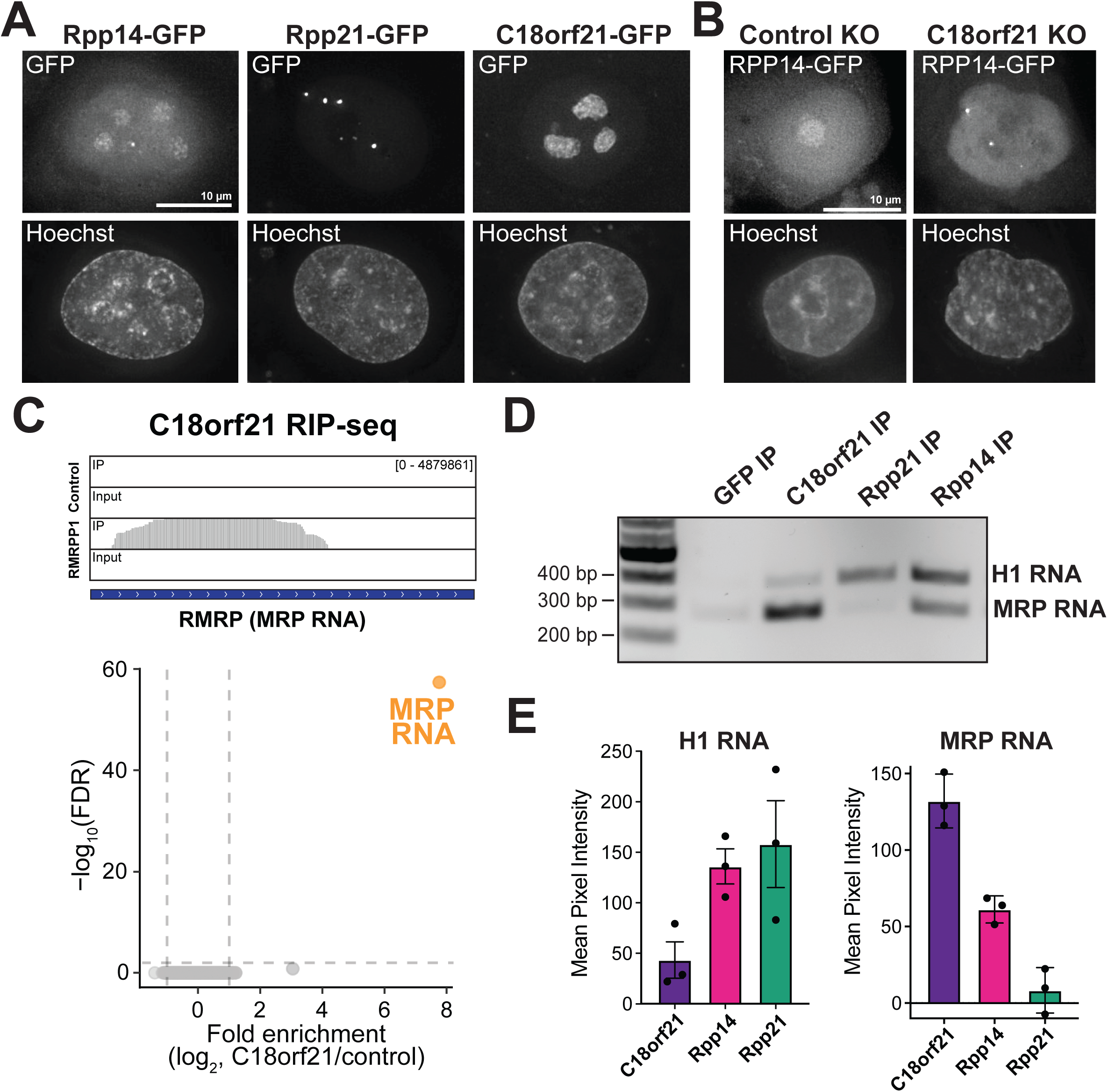
C18orf21 (RMRPP1) is a subunit of the RNase MRP complex. **(A)** Representative Z-projected images taken by live cell imaging cells the ectopically express C-terminally GFP tagged versions of RMRPP1, Rpp21, or Rpp14. Images were deconvolved and each set of images is scaled differently to highlight localization of each component. **(B)** Representative Z-projected images taken by live imaging cells that ectopically express C-terminally GFP tagged Rpp14 and are targeted with either a control guide (AAVS1 locus) or a guide targeting RMRPP1. Images were deconvolved and each image is scaled differently to highlight the differences in localization that were observed. **(C)** Top: RNA-sequencing read coverage plots that display the RMRP reads that were enriched in the RMRPP1 immunoprecipitation. Bottom: A graph showing the Log_10_ False discovery rate (FDR) on the Y-axis and fold enrichment on the X-axis obtained from ribonucleoprotein-immunoprecipitation (RIP) RNA sequencing experiments performed in duplicate. **(D)** Representative agarose gel stained with ethidium bromide showing the results of ribonucleoprotein-immunoprecipitation RT-PCR experiments. **(E)** Quantitation of the mean pixel intensity of either the intensity of the H1 band (left) or RNase MRP RNA band (right) with the GFP RIP RT-PCR background signal subtracted. The error bars represent the standard error of the mean from three replicates.

Although RNase P and MRP share many protein subunits, they have distinct catalytic RNA components (H1 vs. RMRP). Since C18orf21 and Rpp21 display differential localization and form mutually exclusive complexes with the shared subunits of the RNase P and RNase MRP complexes, we sought to determine which RNA C18orf21 associates with in cells. To test whether C18orf21 interacts with H1 RNA or RNase MRP RNA in cells, we performed GFP immunoprecipitations followed by RNA sequencing and RT-PCR (ribonucleoprotein immunoprecipitation (RIP) analysis). We found that immunoprecipitation of Rpp14-GFP isolated both H1 RNA and RNase MRP RNA, consistent with it being a member of both the RNase P and RNase MRP complexes (Fig.2C-E and Extended Data Fig.2A,C-D). C18orf21-GFP isolated the RNase MRP RNA, but not the H1 RNA, whereas Rpp21-GFP pulled down H1 RNA (Fig.2C-E and Extended Data Fig.2B-D). Based on the protein and RNA interactions, genetic associations, and subcellular localizations, we propose that C18orf21 is a specific subunit of RNase MRP and will refer to C18orf21 as RNase MRP Protein 1 (RMRPP1).

### RMRPP1 is structurally-related to Rpp21

To determine the basis for the interaction of RMRPP1 with subunits of the RNase MRP complex, we next analyzed its predicted structural features using Alphafold^39^. Alphafold confidently predicted that the N-terminal domain (residues 1-127) of RMRPP1 folds into an elongated structure with three anti-parallel beta strands sandwiched between two sets of alpha helices, whereas the RMRPP1 C-terminal domain is predicted to be largely unstructured (Fig.3A and Extended Data Fig.3A). AlphaFold predictions of RMRPP1 structures from a variety of organisms suggest that this N-terminal domain is structurally conserved even in cases with low sequence similarity (Extended Data Fig. 3B). Since our data suggests that RMRPP1 is a member of the RNase MRP complex, we used Alphafold 3 to predict how RMRPP1 fits into the larger enzyme complex (Fig. 3B). In this structural model, RMRPP1 is predicted to be sandwiched between Rpp29 and Rpp38, with the two N-terminal helices of RMRPP1 forming a protein-protein interface with Rpp29 (Fig. 3B). To test this structural model directly, we tested if RMRPP1 interacts with Rpp29 in vitro. When expressed on its own in bacteria, the RMRPP1 protein was insoluble. However, upon co-expression of RMRPP1 and Rpp29 in bacteria, we were able to isolate a stable RMRPP1-Rpp29 complex illustrating a direct interaction between RMRPP1 and a component of the RNase MRP complex (Fig.3C).

**Figure 3.**
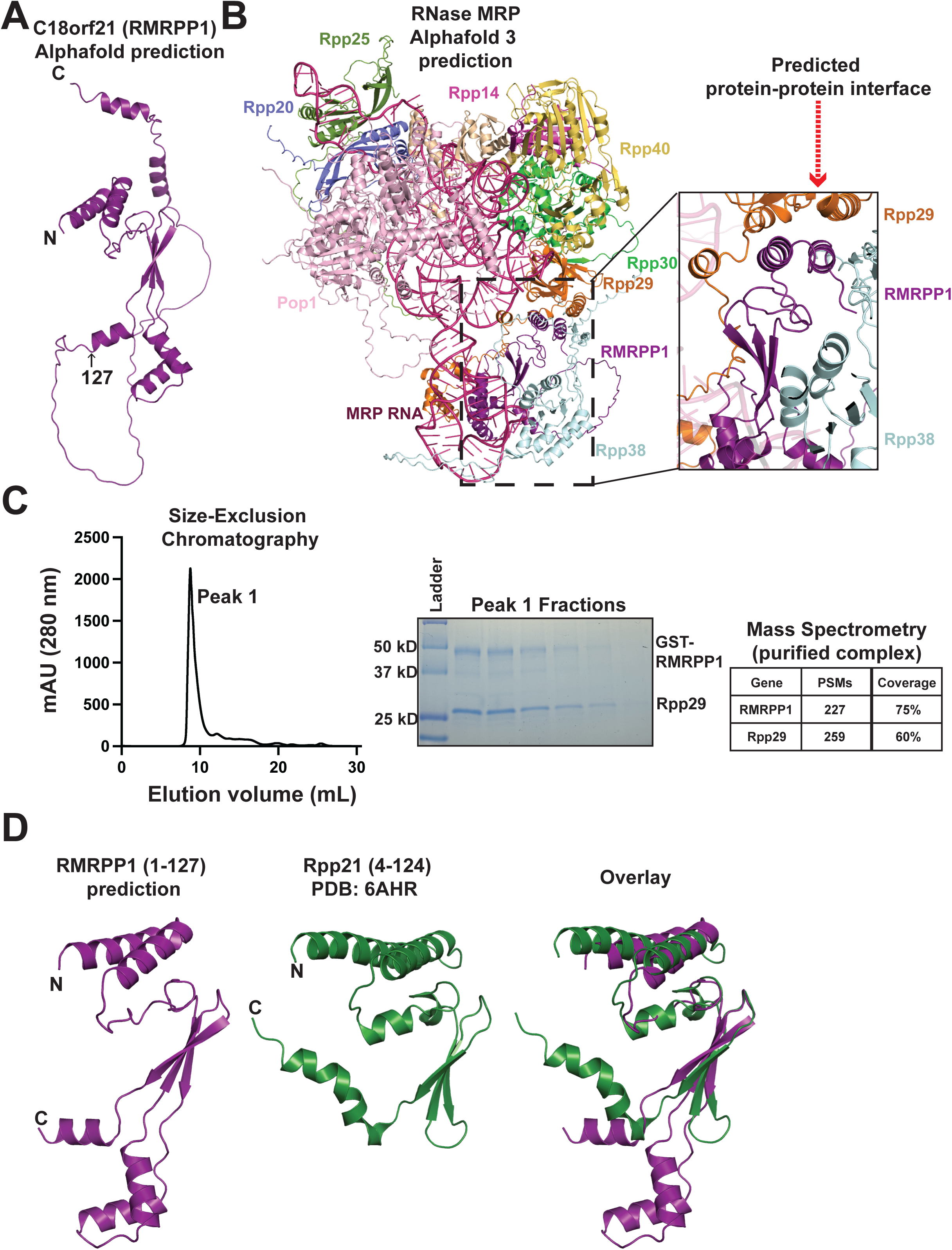
RMRPP1 is predicted to have structural homology with Rpp21. **(A)** Cartoon representation of the structure of RMRPP1 (deep purple) predicted by Alphafold. **(B)** A cartoon representation of the predicted structure of human RNase MRP that was created using AlphaFold 3^39^. Pop5 is shown in wheat, Pop1 is shown in light pink, Rpp40 is shown in yellow orange, Rpp38 is shown in light blue, Rpp30 is shown in green, Rpp29 is shown in orange, Rpp25 is shown in split pea green, Rpp20 is shown in slate blue, Rpp14 is shown in magenta, RMRPP1 is shown in deep purple and MRP RNA is shown in hot pink. The magnified view to the right of the full structure highlights the predicted contacts between RMRPP1 and Rpp29. **(C)** A representative chromatogram resulting from passing the RMRPP1-Rpp29 complex over a Superdex S200-increase column. A coomassie stained SDS-PAGE gel that shows the peak fractions eluted from the Superdex S200-increase column. A table showing the peptide spectrum matches (PSMs) and coverage of RMRPP1 and Rpp29 obtained from injecting the purified complex on the LC-MS. **(D)** A cartoon representation of the predicted structure of RMRPP1 (residues 1-127, deep purple), Rpp21 (residues 4-124, forest green) from PDB ID: 6AHR, and an overlay of the two structures^45^.

Inspection of the previously solved structure of human RNase P shows that the RNase P specific protein Rpp21 similarly uses its two N-terminal helices to interact directly with Rpp29 (Extended Data Fig.3C). Alignment of the structure of Rpp21 and the predicted structure of RMRPP1 revealed that the N-terminal domains of RMRPP1 (residues 1-127) and Rpp21 (4-124) share many structural features despite the fact that RMRPP1 and Rpp21 display only 20% sequence identity (Fig.3D and Extended Data Fig.3D). Each protein has a pair of N-terminal helices that form a hydrophobic surface suggesting each protein interacts with Rpp29 in a similar manner. In addition to the similar N-terminal alpha-helices that are present in both proteins, Rpp21 and RMRPP1 both have three anti-parallel beta-sheets at their N-termini. Specifically, residues 1-61 and 98-115 of the predicted RMRPP1 structure closely matched the structure of residues 19-108 of Rpp21 (Fig.3D and Extended Data Fig.3E). Thus, our analysis suggests that RMRPP1 is related to Rpp21 and that these two proteins act as mutually exclusive subunits of the RNase MRP/P complexes.

### Specificity for the RNase MRP or RNase P complex is dictated by the N-terminus of RMRPP1 and Rpp21

To understand the molecular determinants that allow Rpp21 and RMRPP1 to share structural features, but act as mutually exclusive subunits of the RNase P and RNase MRP complexes, we generated HeLa cell lines that ectopically expressed C-terminally GFP tagged chimeras of Rpp21 or RMRPP1. Since the major differences in the predicted secondary structure of the two proteins are present in their C-terminal domains, we first generated chimeric proteins in which these regions were swapped between Rpp21 and RMRPP1 (Fig.4A). In RIP-RT PCR experiments, we found that the Rpp21^N-term^+RMRPP1^C-term^ construct isolated H1 RNA whereas the RMRPP1^N-term^+Rpp21^C-term^ construct interacted with RNase MRP RNA (Fig.4B). Thus, despite the predicted similarities in the Rpp21 and RMRPP1 structures, these results suggest that the N-terminal domains likely confer specificity for RNase P vs. MRP. To define regions in the N-terminal domain of RMRPP1 that confer RNase MRP complex specificity, we tested Rpp21-RMRPP1 chimeras in which a structurally homologous feature was swapped from RMRPP1 onto Rpp21 (Fig.4C,D and Extended Data Fig.4A). Surprisingly, we found that all three RMRPP1 structural features, when swapped into Rpp21, immunoprecipitated both H1 and MRP RNA (Fig. 4E). Strikingly, the Rpp21^RMRPP1^ ^β-sheets^ construct pulled down more MRP RNA than H1 RNA, similar to that of wild type RMRPP1, suggesting that this region in RMRPP1 is sufficient to confer specificity for MRP RNA (Fig.4E). Together, these results show that substituting any one of the structural features in the N-terminal domain of Rpp21 with the corresponding structural feature of RMRPP1 can confer interaction with RNase MRP RNA.

**Figure 4.**
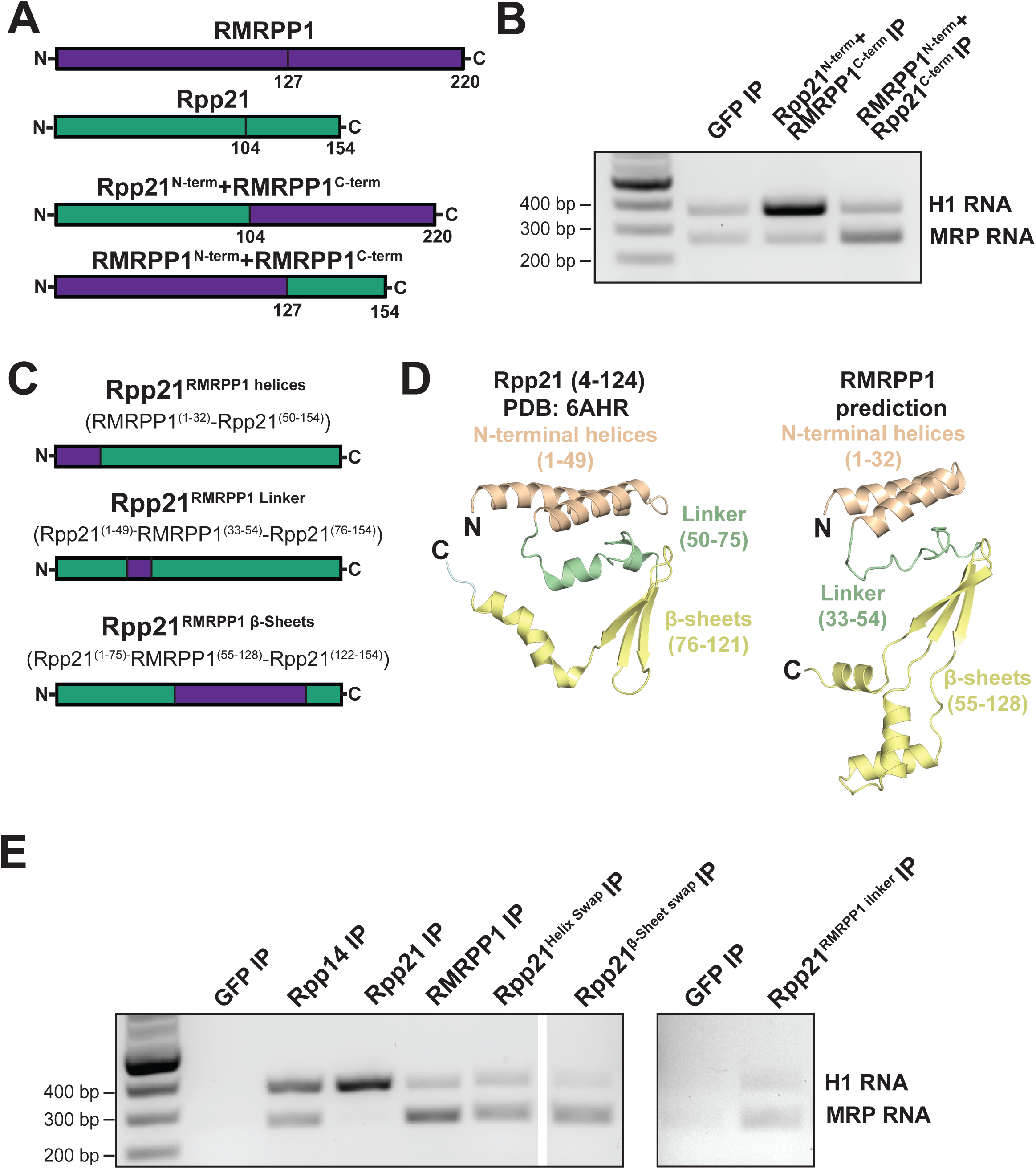
Specificity for RNase MRP or RNase P are dictated by the N-termini of Rpp21 and RMRPP1. **(A)** Diagram depicting the chimeric constructs used in B. **(B)** Representative agarose gel stained with ethidium bromide showing the results of ribonucleoprotein-immunoprecipitation RT-PCR experiments. **(C)** Diagram depicting the chimeric constructs used in D. **(D)** A cartoon representation of the solved structure of human Rpp21 PDB ID: (6AHR) and the predicted structure of RMRPP1. Each of the three shared structural features are depicted in a different color. The N-terminal helices are shown in wheat, the linker is shown in pale green, and the beta-sheets are shown in yellow. **(E)** Representative agarose gel stained with ethidium bromide showing the results of ribonucleoprotein-immunoprecipitation RT-PCR experiments.

### RNase MRP plays an essential and specific role in the biogenesis of the 40S ribosome

The identification of RMRPP1 as an RNase MRP subunit creates the opportunity to assess the functional contributions of RNase MRP. The precise function of RNase MRP in human cells has been difficult to define due to the lack of a specific protein component. Prior work relied on using transient knockout of the RNase MRP RNA to assess of RNase MRP function that relies on a low penetrance dual guide system ^8^. To define the functional contributions of RMRPP1, we tested the consequences of eliminating the gene to cell growth, translation, ribosome biogenesis, and rRNA processing. Based on a competitive growth assay using a co-culture system (see methods), we found that CRISPR/Cas9-based targeting of RMRPP1 led to a substantial reduction in cellular fitness relative to control cells (Fig.5A). This suggests that RMRPP1 is required for cellular viability consistent with prior work that the RNase MRP RNA is essential in yeast and human cells^8,21^.

**Figure 5.**
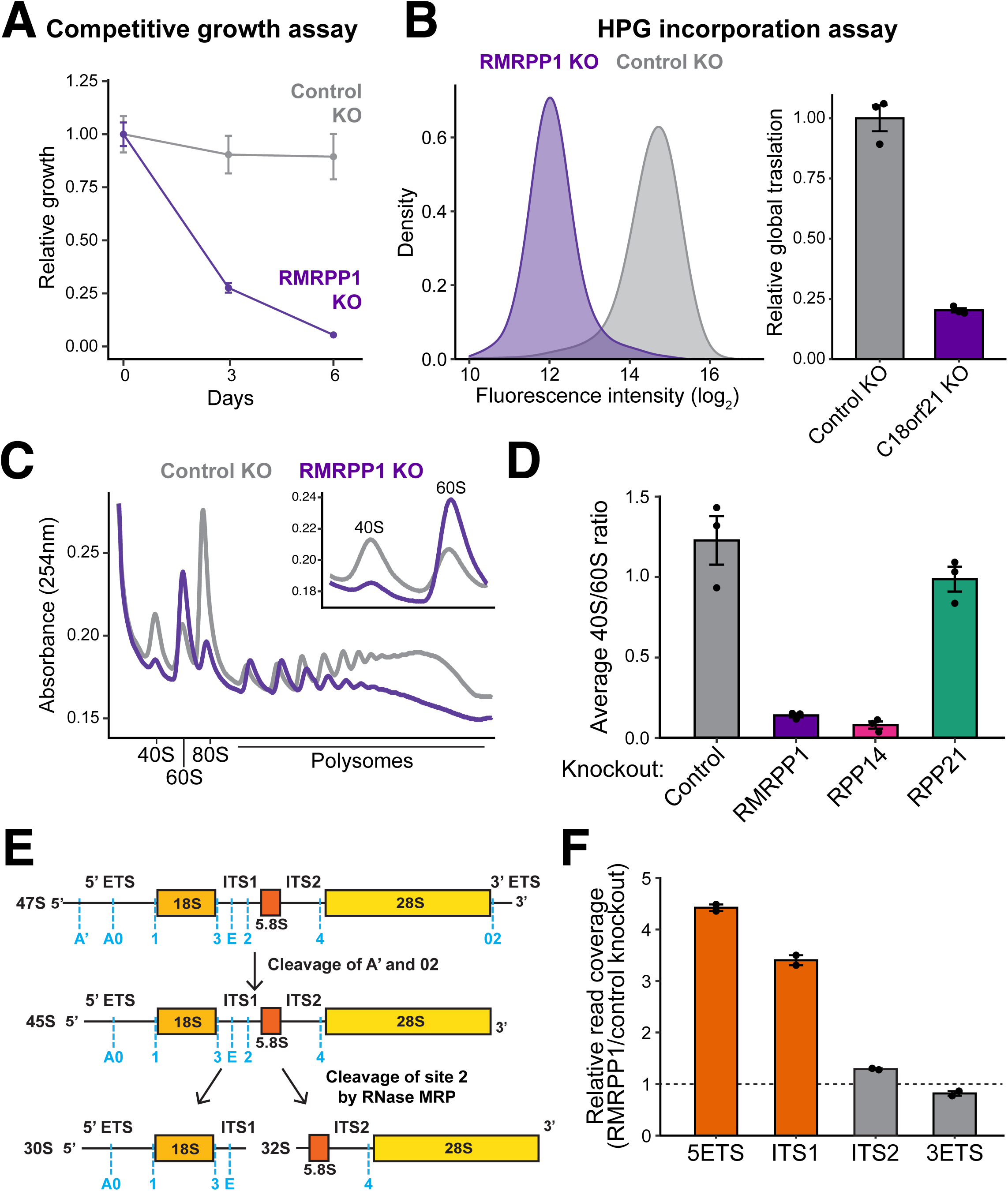
Loss of RMRPP1 leads to defects in ribosome biogenesis. **(A)** A plot showing the relative growth (Y-axis) of either RMRPP1 knockout cells (purple) or control knockout cells (grey) relative to control cells over the course of time (X-axis). **(B)** A plot generated by measuring the fluorescence intensity (X-axis) of HPG incorporated cells by flow cytometry (Left). Quantitation of average HPG signal in each condition. The error bars represent standard error of the mean from three replicates. **(C)** An overlay of the polysome profiles obtained from the fractionation of cell lysates separated on a 10 – 50% sucrose gradient. The inset on the top right shows a magnified view of the 40S and 60S traces. **(D)** Quantification of the ratio of 40S to 60S ribosomes obtained from polysome gradients with the indicated gene knocked out by CRISPR-Cas9 targeting. The error bars represent the standard error of the mean from three biological replicates. (**E**) Schematic outline of the primary rRNA maturation pathway in HeLa cells^3^. **(F)** Quantitation of RNA sequencing data showing the relative read coverage of different domains of the 45S rRNA transcript obtained from RNA isolated from RMRPP1 knockout cells and AAVS1 control knockout cells. The error bars represent the standard error of the mean from 2 replicates.

As RNase MRP is predicted to contribute to ribosome production, we compared rates of protein synthesis using a L-Homopropargylglycine (HPG, a clickable methionine analog^40^) incorporation assay. RMRPP1 knock out cells displayed substantially reduced global translation rates compared to controls (Fig.5B). Consistent with the HPG incorporation assays, RMRPP1 knockout HeLa cells also displayed a decreased polysome/monosome ratio compared to control knockout cells (Fig.5C). Strikingly, we found that the fraction of mature 40S ribosomes was depleted whereas the number of free 60S subunits increased in RMRPP1 knockout cells compared to control HeLa cells (Fig.5C,D). In addition, we found that the knockout of RPP14, but not RPP21, resulted in a similar decrease in 40/60S ribosome ratio (Fig.5D and Extended Data Fig.5A,B). Thus, this unexpected defect in the ratio of mature ribosome species is specific to eliminating RNase MRP activity.

The cleavage activity of RNase MRP separates the 18S rRNA subunit from the 5.8S and 28S rRNA subunits facilitating the maturation of each rRNA subunit (Fig.5E). To test if the changes in ribosomal subunit population were due to changes in rRNA processing, we compared the abundance of mature rRNA species in control cells and in Rpp21, RMRPP1, and Rpp14 knockout cells. Due to its RNase P-specific role, we found that there was no difference in mature rRNA species between control and Rpp21 knockout cells (Extended Data Fig.5C). Consistent with a role in RNase MRP complex function, we found that RMRPP1 and Rpp14 knockout cells each displayed decreased amounts of mature steady state 18S relative to 28S rRNA (Extended Data Fig.5C,D). The analysis of RMRPP1 knockout cells is consistent with prior work showing that that targeting RNase MRP RNA with CRISPR-Cas9 leads to an increase in the abundance of rRNA processing intermediates that correspond to changes in ITS1 cleavage^8^. Knockout of RMRPP1 also resulted in the increase in rRNA intermediates, particularly upstream of the 5.8S rRNA (Fig.5F). Previous work suggests that RNase MRP cleavage at site 2 in ITS1 is a step in the predominant pathway in HeLa cells for the production of 18S, 5.8S, and 28S rRNA (Fig.5E) ^3^. Surprisingly, we found that RNase MRP is only essential for 18S rRNA production (Fig.5C). This suggests the existence of an alternative rRNA processing pathway that bypasses loss of RNase MRP to generate 5.8S and 28S rRNA.

In addition to rRNA level changes, we also assessed the transcriptome of cells lacking RNase MRP activity (RMRPP1 knockout) in HeLa cells. RMRPP1 knockouts revealed strong changes in hundreds of mRNAs (Extended Data Fig. 5E). These included upregulation of several rRNA processing components, such as UTP4, WDR75, WDR43, which may represent compensatory mechanisms upon loss of RNase MRP activity (Extended Data Fig. 5E). We also observed hundreds of mRNAs downregulated in RMRPP1 knockout, including downregulation interferon and complement system genes (Extended Data Fig. 5E,F). These gene expression changes in RNase MRP deficient cells may be related to how RNase MRP diseases are associated with immunodeficiencies. Together, this functional analysis indicates that the RNase MRP-specific protein RMRPP1 is essential for cell viability and has key roles translation, 40S ribosome biogenesis, and provides insights on how loss of RNase MRP activity alters the transcriptome.

### Nepro is a multi-functional protein that associates with the RNase MRP complex

The discovery of RMRPP1 as a unique member of the RNase MRP complex allows for the identification of additional proteins that interact with the RNase MRP complex, but not the RNase P complex. To evaluate if there are any such proteins, we compared the proteins that interact with Rpp21 and RMRPP1 in our IP-MS experiments. The protein Nepro (C3orf17) stands out as a unique interacting partner for both RMRPP1 and Rpp14 but not Rpp21 (Fig. 6A and Extended Data Fig.6A). Based on the Depmap database, Nepro displays genetic correlations with several of the shared RNase P/MRP complex components and the RNase MRP specific protein RMRPP1 (Fig.6B). Similarly, Nepro clusters with factors involved in translation in both genome-wide Perturb-Seq and a pooled optical CRISPR-Cas9 based functional screen^37,41^.

**Figure 6.**
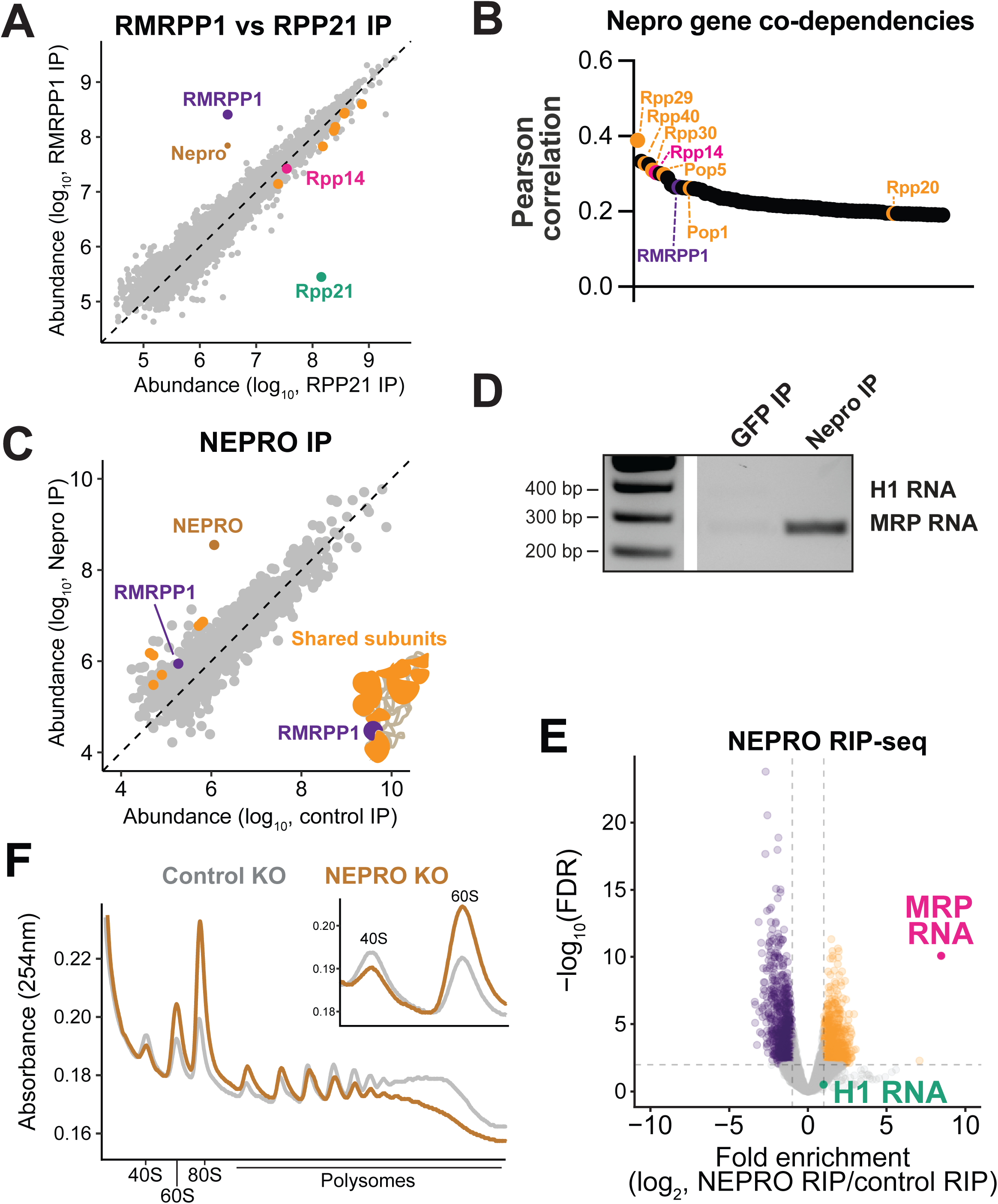
Nepro is an rRNA processing factor that associates with RNase MRP. **(A)** A Scatter plot showing the abundance of the proteins detected in the indicated immunoprecipitation-mass spectrometry experiments. The shared components of the RNase P/MRP complexes are highlighted in orange, except Rpp14 which is shown in pink. RMRPP1 is highlighted in purple, Nepro is highlighted in brown, and Rpp21 is highlighted in green. **(B)** A waterfall plot showing the genes with the highest Pearson correlation of CRISPR-Cas9 based targeting effect scores to Nepro in the Depmap database. The shared RNase P and MRP components are shown in orange, except Rpp14 is shown in pink, and RMRPP1 is shown in purple. **(C)** A Scatter plot showing the abundance of the proteins detected in the indicated immunoprecipitation-mass spectrometry experiments. The shared components of the RNase P/MRP complexes are highlighted in orange. RMRPP1 is highlighted in purple, and Nepro is highlighted in brown. Peptides mapping to Rpp21 were not identified in the Nepro IP-MS experiments. **(D)** Representative agarose gel stained with ethidium bromide showing the results of ribonucleoprotein-immunoprecipitation RT-PCR experiments performed in triplicate. **(E)** A graph showing the Log_10_ False discovery rate (FDR) on the Y-axis and fold enrichment on the X-axis obtained from ribonucleoprotein-immunoprecipitation (RIP) RNA sequencing experiments. **(F)** An overlay of the polysome profiles obtained from the fractionation of cell lysates separated on a 10 – 50% sucrose gradient. The Nepro profile is depicted in brown and the control profile is depicted in gray. The inset on the top right shows a magnified view of the 40S and 60S traces.

To determine the interacting partners of Nepro, we conducted immunoprecipitation-mass spectrometry experiments in HeLa cells ectopically expressing a C-terminally tagged Nepro-GFP fusion. We found that Nepro interacts with the shared components of the RNase P/MRP complex and the RNase MRP specific protein RMRPP1, but not the RNase P specific protein Rpp21 (Fig.6C). This observation is supported by previous work that showed Nepro can interact with several of the shared RNase P/MRP components^42^. To determine if Nepro has specificity for either H1 RNA or RNase MRP RNA, we performed GFP immunoprecipitations followed by RNA sequencing and RT-PCR. Nepro-GFP immunoprecipitation isolated RNase MRP RNA but not H1 RNA, suggesting that Nepro associates with the RNase MRP complex, but not the RNase P complex (Fig.6D-E and Extended Data Fig.6B). To test if loss of Nepro leads to a similar defect in ribosome biogenesis as loss of RMRPP1 or Rpp14, we carried out polysome profiling in HeLa cells in which Nepro was targeted by CRISPR-Cas9. Upon Nepro depletion, we observed a decrease in the population of mature 40S ribosomes that mirrored the decrease observed when RMRPP1 or Rpp14 was knocked out (Fig.6F). Together this data shows that Nepro associates with RNase MRP complex and has a key role in ribosome biogenesis.

Previous work highlighted multiple additional roles for Nepro across different cell types. For example, Nepro is activated downstream of Notch signaling to inhibit differentiation of neurons, and loss of Nepro leads to the mislocalization of ribosomal subunits and mitochondrial proteins at the two-cell stage in mouse embryos^43,44^. In agreement with roles for Nepro outside of the RNase MRP complex, we found that Nepro interacts with other rRNA processing factors in our IP-MS experiments that do not immunoprecipitate with RMRPP1 or Rpp14 (Extended Data Fig.6C). Additionally, we found that Nepro pulls down several proteins that are associated with genome organization and integrity (Extended Data Fig.6C). Similarly, using RIP-Seq we found that RMRPP1, Rpp14, and Nepro interact with RNase MRP RNA. However, in contrast to RMRPP1 and Rpp14, we found that Nepro significantly interacts with 875 RNAs, including the many histone H1 subtypes, proteasome subunits, and nuclear-encoded mitochondrial mRNAs highlighting potential roles outside of rRNA processing for this protein (Fig. 6D and Extended Data Fig.6D). Together our work indicates that Nepro associates specifically with the RNase MRP complex, but not RNase P. However, our results also suggest that RMRPP1 is the only RNase MRP specific protein as Nepro additionally associates with unrelated complexes.

## Discussion

Despite the distinct roles of RNase MRP and RNase P in cellular function, they share multiple protein components and it was unknown how the function of RNase MRP is specified from RNase P at a protein level. To address this, we identified the previously uncharacterized protein C18orf21/RMRPP1 as an RNase MRP-specific protein that is conserved across vertebrates. Our work showed that loss of RMRPP1 leads to defects in cellular fitness, ribosome biogenesis, and rRNA maturation highlighting its central role in RNase MRP function. The finding of RMRPP1 as an RNase MRP specific protein led us to identify Nepro as an additional interactor of the RNase MRP complex, but not RNase P. However, unlike RMRPP1, Nepro has several additional protein and RNA interaction partners outside of the RNase MRP complex, highlighting multiple roles for the protein outside of ribosome biogenesis, placing RMRPP1 as the only unique interacting partner of the RNase MRP complex.

Our understanding of the mechanisms that RNase P uses to recognize and process tRNA is informed by several structural studies of the enzyme across all three domains of life^30,45–47^. In contrast, most of the molecular information about the enzymatic action of RNase MRP comes from studies of the enzyme from *S. cerevisiae*^32,33^. For example, the structure of *S. cerevisiae* RNase MRP bound to a 21 base pair oligomer composed of part of the ITS1 sequence shed light on the catalytic mechanism the enzyme uses to cleave single-stranded RNA. However, no structural or biochemical experiments have identified how RNase MRP recognizes its substrate^12^. The studies of yeast RNase MRP have also shed light on the overall organization of the enzyme and defined how two RNase MRP specific proteins found in *S. cerevisiae*, Snm1 and Rmp1, fit into the complex^32,33^. Strikingly, Snm1 and Rmp1 both have several structural elements that match those found in our prediction of RMRPP1 and Nepro, but also have regions that diverge significantly (Extended Data Fig.6E). For example, Rmp1 forms a bundle of helices at its N-terminus that interact with both Pop1 and the RNase MRP RNA subunit (Nme1)^11,12^. AlphaFold 3 predicts that Nepro has a similar bundle of helices at its N-terminus as Rmp1, but Nepro has an additional 366 residues at its C-terminus (Extended Data Fig.6E). Although the structures of *S. cerevisiae* RNase MRP provide important models of the complex, several open questions remain about the differences between the yeast and human RNase MRP specific proteins, including how the human proteins fit into the enzyme. Ultimately, the identification of RMRPP1 and Nepro open new paths to answer open questions surrounding the architecture and substrate specificity of the human RNase MRP complex.

Previous work found that human RNase MRP is responsible for cleaving site 2 in ITS1 of pre-rRNA to separate the 18S rRNA from the 5.8S and 28S rRNA subunits (Fig.5E)^8^. We found that loss of RMRPP1 resulted in the depletion of mature 18S rRNA species and an increase in sequencing reads in both the 5’ ETS and ITS1, indicating a buildup of rRNA processing intermediates (Fig.5F). Similarly, studies conducted in patient-derived fibroblast cells with a mutation in RNase MRP RNA (RMRP 70^AG^) found no change in 5.8S rRNA maturation, but found an increase in the abundance of intermediate 18S rRNA species^48^. Together with previous studies, our work demonstrates that the maturation of the 18S rRNA relies strongly on RNase MRP function^8,48^. This could be through cleavage at site 2 in ITS1, but it is also possible the RNase MRP cleaves at a second location that is required for 18S rRNA biogenesis.

In agreement with our observed rRNA processing defects, our polysome profiling experiments show a depletion of the 40S ribosome and a buildup of the 60S ribosome upon RMRPP1, Nepro, and Rpp14 knockout (Fig.5C,D). Our results strongly suggest that RNase MRP is dispensable for the production of 60S ribosomes, consistent with the genetic interaction between RMRPP1 and 40S ribosome biogenesis factors^37^. However, prior work suggested that RNase MRP is important for the production of 5.8S and 28S rRNA^3^. The buildup of the 60S ribosome in our experiments suggests the existence of an alternative pathway that acts to ensure proper 5.8S and 28S rRNA biogenesis in the absence of RNase MRP activity. It is possible this could include an unidentified redundant or alternative pathway that allows for the correct processing of the 5.8S subunit, but not the 18S rRNA. Together, our studies have leveraged the discovery of a RNase MRP-specific protein to define the contributions of this enzyme to rRNA processing and ribosome biogenesis in cells.

## Acknowledgements

We thank members of the Cheeseman laboratory for discussions and assistance. and David Bartel’s laboratory for use of equipment and reagents. Research was supported by grants from the NIH/National Institute of General Medical Sciences (R35GM126930) and the Chan Zuckerberg Initiative to I.M.C., an American Cancer Society Postdoctoral Fellowship (134255-PF-20-041-01-DMC) to E.M.S. and a graduate fellowship from the Natural Sciences and Engineering Research Council of Canada Postgraduate Fellowship to J.L. We also thank the Whitehead Institute Quantitative Proteomics, Genome technology, and Flow Cytometry cores for their help and support.

## Declaration of interests

The authors declare no competing interests.

## Contributions

Conceptualization: JL, EMS, IMC; Methodology: EMS and JL; Investigation: EMS, JL, SH; Formal analysis: EMS and JL; Writing: EMS, JL, and IMC; Supervision: IMC; Funding acquisition: IMC, EMS, and JL

## Experimental Procedures

### Plasmid Constructs

The bicistronic cassette used to co-express C18orf21 and Rpp29 in *E. coli* was synthesized by Twist Bioscience. Restriction cloning was used to insert the cassette into the pGEX-6P-1 vector. Single guide RNAs (sgRNA) were synthesized as complimentary ssDNA pairs by Integrated DNA Technologies (IDT) that were annealed before being inserted into the LentiCRISPRv2-Opti vector (Addgene: 163126). C18orf21, and Nepro resistant to the corresponding sgRNA was synthesized by Twist Bioscience. Rpp21, and Rpp14 genes were amplified from cDNA obtained from HeLa cells.

### Lentiviral Production and transduction

Lentivirus was produced by transfecting HEK293T cells with 1.2 µg lentiCas9v2-Opti vector containing a sgRNA, 1 µg psPAX2 (Addgene: 12260) and 0.4 µg vsFULL (Gift from Sabatini lab) using Xtremegene-9 (Roche). Approximately 14-16 hours after transfection the growth medium was changed. 24 hours after changing media the lentivirus was harvested and stored at −80 °C.

The resulting sgRNA-Cas9 lentivirus was used to transduce HeLa cells with 10 μg/ml polybrene and spinfection at 2250x g for 30 minutes at 37 °C. The cells were incubated for 16 hours. The media was replaced and then cells were grown for an additional 24 hours. The infected cells were selected for using 0.4 μg/ml puromycin for 48 hours. The selected cells were grown for 3 days prior to analysis of global translation, polysome profile, and rRNA and mRNA species.

### Tissue Culture

Except for HPG incorporation experiments, all cells were cultured at 37 °C with 5% CO2 in Dulbecco’s modified Eagle medium supplemented with 10% tetracycline-free fetal bovine serum, 100 U/ml penicillin-streptomycin, and 2 mM L-glutamine. Cells were counted using a Z2 Coulter counter (Beckman Coulter).

### Cell Line generation

Cell lines that have doxycycline inducible expression of C-terminally GFP tagged C18orf21, Nepro, Rpp21, or Rpp14 were created as follows. First, Gibson assembly was used to insert each construct downstream of a tetracycline-responsive promoter and upstream of Myc – TEV – GFP. These elements were flanked on either side by ∼800 bp homology arms for insertion into the AAVS1 safe harbor locus and contained a puromycin resistance marker ^34^. Each cell line was created by transiently transfecting this donor plasmid along with the pX330 plasmid containing a sgRNA targeting the AAVS1 locus using Lipofectamine 2000 transfection reagent (Invitrogen), according to the manufacturer’s instructions. Cells were selected with puromycin (0.4 μg/ml) for 4 days.

### Competitive growth assays

To determine if RMRPP1 is essential for cellular fitness, we used a competitive growth assay in which control HeLa cells (labeled with mCherry) were co-cultured with cells where RMRPP1 was knocked out by CRISPR-Cas9 (unlabeled cells) allowing us to monitor the relative growth of the cells grown together by flow cytometry. Specifically, HeLa cells were spinfected with lentivirus that contained Cas9 and C18orf21 or control AAVS1 sgRNA. Infected cells were selected with puromycin for 2 days. After selection, the infected cells were mixed 50:50 with mCherry expressing HeLa cells and the relative ratio of mCherry/uncolored cells were monitored every three days using BD FACSSymphony A1 Cell Analyzer (BD Biosciences).

### Live Cell Imaging

For live cell imaging experiments 50,000 cells were plated into the wells of a 12 well polymer glass bottom plate (Cellvis). Expression of each GFP construct was induced by the addition of doxycycline (1 μg/ml) for 48 hours. Hoechst (0.1ug/ml) was added to each well 1 hour before imaging. To perform the Rpp14 imaging experiments with C18orf21 knockout or a control knockout the following protocol was followed. On the first day of the experiment the HeLa cell line with the GFP tagged Rpp14 construct under doxycycline inducible expression were plated at ∼40% confluency (0.5 x 10^6^ cells) in a 6 well dish. The following day expression of Rpp14-GFP was induced by addition of doxycycline and cells were transduced with a lentivirus that contained Cas9 and a sgRNA that targeted either C18orf21 or AAVS locus in the genome. Additionally, the virus contained a puromycin resistance gene that allows for the selection of infected cells. Puromycin selection was carried out for three days and on the third day ∼50,000 surviving cells were transferred to a 12 well polymer bottom plate to allow for live cell imaging. Images were taken ∼24 hours after plating. All live cell imaging was performed with a DeltaVision Ultra High-Resolution microscope with a 60x/1.42NA objective (Cytiva). ImageJ was used to perform all image analysis and manipulation.

### Protein Purification

To express and purify C18orf21-Rpp29 the following protocol was followed. *Escherichia coli* BL21-DE3 cells were grown at 37 °C until they reached an OD_600_ of 0.6-0.8. The cells were induced with 250 mM Isopropyl β-d-1-thiogalactopyranoside (IPTG) for 16 hours at 18 °C. Cells were pelleted at 4,000 x g in a JLA8.1000 (Beckman Coulter) in a Avanti JXN-26 centrifuge (Beckman Coulter) before being flash frozen in liquid nitrogen and stored at −80 °C. Cells were thawed in a 25 °C water bath and then subsequently resuspended in lysis buffer (50 mM Tris HCl pH 7.5, 200 mM NaCl, 5 mM beta-mercaptoethanol (BME), 1 mM phenylmethylsulfonyl fluoride (PMSF) supplemented with 1 cOmplete protease inhibitor cocktail tablet (Roche, 04693159001). After resuspension cells were lysed by sonication (10 seconds on, 30 seconds off for 2.5 minutes). The lysate was clarified by centrifugation at 18,000 rpm for 1 hour at 4 °C in a JA-17 rotor (Beckman Coulter) in an Avanti JE centrifuge Beckman Coulter). Clarified lysate was passed over a Glutathione-Agarose (GE Healthcare) column that was equilibrated with lysis buffer. The column was then washed 20 column volumes of 50 mM Trish HCL pH 7.5, 200 mM NaCl, 1 mM DTT. To elute the protein beads were nutated with precision protease overnight at 4 °C or 5 column volumes of 25 mM Tris HCl pH 8, 50 mM NaCl, 1 mM dithiothreitol (DTT), 50 mM reduced glutathione was used. Eluted protein was dialyzed into S200 buffer (25 mM Tris HCl pH 8, 100 mM NaCl, 1 mM dithiothreitol (DTT)) at 4 °C overnight. Dialyzed protein was concentrated and passed over a S200 increase column 10/300 GL (Cytiva). Peak fractions were analyzed by Coomassie stained SDS-PAGE gels to determine purity. Pure fractions were then pooled, concentrated, flash frozen in liquid nitrogen and stored at −80 °C.

### Immunoprecipitation-mass spectrometry

Expression of each GFP construct was induced by the addition of doxycycline for 48 hours. After dox induction cells were harvested after incubation with 5 mM EDTA in 1X PBS for 5 minutes at 37 °C. Harvested cells were pelleted by centrifugation at 200 x g. Cells were washed once with 1X PBS and then once with 1x lysis buffer (25 mM HEPES pH 8.0, 2 mM MgCl2, 0.1 mM EDTA pH 8.0, 0.5 mM EGTA pH 8.0, 300 mM KCl and 10% glycerol). Cells were pelleted between each wash step. Finally, the cells were resuspended in a 1:1 ratio in 1x lysis buffer and flash frozen in liquid nitrogen before being stored at −80 °C. Polysome lysis buffer (20 mM Hepes pH 7.5, 100 mM KCl, 5 mM MgCl_2_, 1% Triton X-100) supplemented with 1% CHAPS, 1 cOmplete EDTA-free protease inhibitor cocktail (Roche), and 0.02 U/μl of SUPERaseIn (Invitrogen) was added to each frozen cell pellet before thawing the cells in a 37 °C water bath. Cells were lysed by 10 cycles of sonication at high amplitude with a 30 second on time and 1 minute off time (Bioruptor, Diagenode). The lysate was clarified by centrifugation at 21,000 x g for 30 minutes at 4 °C. Clarified lysate was then incubated with Protein A beads (BioRad) coupled to a rabbit anti-GFP antibody ^49^ at 4 °C for 2 hours on a rotating wheel. Following incubation with the clarified lysate the beads were washed 4 times for 5 minutes with polysome lysis buffer supplemented with 1 mM DTT, 10 µg/ml leupeptin/pepstatin/chymostatin, and 0.02 U/μl of SUPERaseIn. Three elutions were performed by rotating the beads with 2 bead volumes of 100 mM glycine pH 2.6 for 5 minutes at 4 °C. The three elutions were pooled and Tris pH 8.5 was added to a final concentration of 200 mM. Following elution 1/5 the final volume of trichloroacetic acid was added to each elution. The eluted proteins were precipitated overnight at 4 °C. The precipitated proteins were pelleted by centrifugation at 21,000 x g for 30 minutes. The proteins were then washed three times with 1 ml ice cold acetone before being dried in a speedvac for 5 minutes at 30 °C. The proteins were then resuspended in 5% SDS, 50 mM tetraethylammonium bromide pH 8.5 (TEAB), 20 mM DTT and then were incubated at 95 °C for 10 minutes. The resuspended proteins were then allowed to cool to room temperature. Next, 40 mM iodoacetamide was added to the resuspended proteins and the alkylating reaction was allowed to proceed for 30 minutes in the dark before being quenched by the addition of 2.5% v/v phosphoric acid. Following alkylation 6 times the volume of S-trap binding buffer (90% MeOH, 100 mM TEAB, pH 7.55) was added and the proteins were loaded onto an S-Trap microcolumn (Protifi) by centrifugation at 4000 x g for 1 minute. The eluate was passed through the column a second time and subsequently the column was washed 4 times with S-trap binding buffer. An on-column digestion was performed by adding 1 µg of trypsin diluted in 50 mM TEAB pH 8.5 to the top of the column and placing the column in a humidified chamber at 37 °C overnight. Following overnight digestion, the tryptic peptides were eluted with 40 µl of 50 mM TEAB, followed by 40 µl of 0.2% formic acid, and finally with 35 µl 50% acetonitrile/0.2% formic acid. The eluted peptides were pooled, flash frozen, and then lyophilized overnight. The amount of eluted peptide obtained was quantified using the Pierce fluorometric peptide assay kit (Pierce) following the manufacture’s protocol. The lyophilized peptides were resuspended in 0.1% formic acid to a final concentration of 250 ng/uL. 250 ng of each sample was then loaded onto an Exploris 480 Orbitrap mass spectrometer equipped with a FAIMS Pro source connected to an EASY-nLC chromatography system. The injected peptides were separated on a 25 cm analytical column (PepMap RSLC C18 3 µm, 100A, 75 µm). Liquid chromatography was performed with a flow rate of 300 nl/min. A 120 minute step gradient was performed as follows. Step 1) 41 minutes 6–21% B, Step 2) 21–36% B for 20 min, Step 3) 36–50% B for 10 min, Step 4) 50–100% B over 15 min, Step 5) 100–2% B for 6 minutes, Step 6) 2–100% for 6 minutes. The orbitrap and FAIMS were operated in positive ion mode with a positive ion voltage of 1800 V; with an ion transfer tube temperature of 270 °C; using standard FAIMS resolution and compensation voltages of –45 and –65 V. Full scan spectra were acquired in profile mode at a resolution of 120,000, with a scan range of 350–1200 m/z, automatically determined maximum fill time, standard AGC target, intensity threshold of 5×103, 2–5 charge state, and dynamic exclusion of 30 seconds.

Proteome Discoverer 2.4 (Thermo Fisher Scientific) was used to process the raw files. Sequest HT was used to identify proteins and peptides (Thermo Fisher Scientific). The human protein database (UP000005640) with the sequence of GFP added was used as the reference database. Tryptic peptides with up to 2 missed cleavages were allowed. Mass tolerances for precursors was set to 10 ppm and fragments were set to 0.02 Da. The analysis included the following post-translational modifications: dynamic oxidation (+15.995 Da; M), dynamic acetylation (+42.011 Da; N-terminus), dynamic Met-loss (–131.04 Da; M N-terminus), dynamic Met-loss combined with acetylation (– 89.03 Da; M N-terminus), and static carbamidomethylation (+57.021 Da; C). Percolator was used to filter the identified peptides to ensure a FDR of less than 0.01.

### Ribonucleoprotein-immunoprecipitation

Cells were harvested and immunoprecipitations were carried out as described above in the IP-MS methods section. There were two major changes to the above protocol. First, a GFP nanobody was coupled to NHS Mag Sepharose beads (Cytiva) for the immunoprecipitation and second, instead of eluting the bound proteins off with glycine, TRI reagent (Invitrogen, AM9738) was added to the beads.

### RNA extraction

400uL TRI Reagent (Invitrogen, AM9738) was added directly to beads or cells. RNA was then purified using Phasemaker tubes (Invitrogen, A33248). The purified RNA pellet was washed with 70% ethanol and resuspended in RNase free water. The RNA concentration was quantified by nanodrop. For analysis of RNA integrity and to monitor mature 18/28S levels, the total RNA was diluted to 5 ng/μl and run on an Agilent BioAnalyzer 2100 using the Agilent RNA 6000 Pico Reagents Kit (5067-1513).

### RT-PCR

After isolation of the bound RNA reverse transcription reactions were performed using the Maxima First Strand cDNA synthesis kit (Thermo Scientific) following the manufacturer’s protocol. Following cDNA synthesis, PCR reactions were performed with Q5 2X master mix (New England Biolabs). Primers complementary to both H1 RNA and MRP RNA were added to the reaction at a final concentration of 500 nM each. The forward and reverse H1 RNA primers had 24 additional bases added to their 5’ ends to help separate the H1 PCR product from the MRP PCR product. PCR products were electrophoresed on a 1.5% agarose gel. Bands were visualized by staining the gel with ethidium bromide and the gel was imaged with a Chemidoc Imaging System (Biorad). ImageJ was used to quantify the intensity of each band and Prism (GraphPad) was used to analyze and plot the data.

### RNA sequencing

For RIP-seq library preparation, libraries were prepared with the Watchmaker library prep kit (Watchmaker Genomics) with a QiaFast Select rRNA kit (Qiagen) module for rRNA depletion and each step was performed according to the manufacturer’s instructions. For the analysis of rRNA from knockout cells, the total RNA was directly prepped using the Kapa HyperPrep kit without any depletion/enrichment modules according the manufacturer’s instructions. The libraries were sequenced using the Element AVITI with standard 150×150bp paired end reads.

### Analysis of RNA sequencing

For the RIP-seq experiments, reads were mapped to the genome using STAR with the following options --runThreadN 6 --runMode alignReads --outFilterMultimapNmax 1 --outFilterType BySJout --outSAMattributes All -- outSAMtype BAM SortedByCoordinate. Sequencing reads for knockout cells were mapped to the human 45s rRNA STAR with the following options --runThreadN 2 -- limitBAMsortRAM 10000000000 --runMode alignReads --outFilterMultimapNmax 5000 - -winAnchorMultimapNmax 1000 --outFilterType BySJout --outSAMattributes All -- outSAMtype BAM SortedByCoordinate. For RIP-seq experiments, reads were mapped to the human genomic sequences and annotations were downloaded from the GENCODE website (release 25, GRCh38.p7, primary assembly or main annotation). For knockout experiments, the reads were mapped to a custom genome containing the sequence of the human precursor 45S ribosomal RNA (Gene ID: 100861532. Exon or rRNA mapping reads were quantified using htseq-count (0.11.0) with the parameters -f bam -t exon -s reverse –nonunique all and a read cuttoff of ≥20 reads in each sample was applied for each gene. For rRNA, a custom annotation file was generated containing the human 5’ ETS, 18S, ITS1, 5.8S, ITS2, 28S, and 3’ETS. Statistical analysis of immunoprecipitation fold enrichment was calculated with DESeq2 using the interaction term to normalize IP to input RNA sequencing counts. For quantification of rRNA, the raw HTseq-count reads for each rRNA region were normalized to the total reads mapping to rRNA then compared to the control.

### Homopropargylglycine (HPG) incorporation assays

Prior to HPG incorporation, the media was replaced with methionine free media for 30 minutes. Cells were labelled with 500 μM L-Homopropargylglycine in methionine free media and incubated at 37 °C for 30 minutes. After amino acid incorporation, cells were washed twice with PBS, trypsinized, and fixed with 4% formaldehyde in PBS for 15 minutes at room temperature. Fixed cells were washed once with PBS then blocked using AbDil (20 mM Tris-HCl, 150 mM NaCl, 0.1% Triton X-100, 3% bovine serum albumin, 0.1% NaN3, pH 7.5) for 30 minutes at room temperature. Click chemistry was performed in solution by incubating the fixed cells in 120 μL of 100 mM Tris pH 8.0, 1 mM CuSO4, 5 μM AlexaFluor Azide, and 0.1 M ascorbic acid at room temperature for 30 minutes in the dark. Cells were washed 3 times with PBS containing 0.1% Triton X-100, strained, and analyzed by flow cytometry using the BD FACSSymphony A1 Cell Analyzer (BD Biosciences) and FlowJo. Dead cells and cell doublets were removed from the analysis on the basis of forward and side scatter.

### Polysome gradient analysis

One 15 cm plate cells at ∼80% confluency was treated with 100 μg/mL cycloheximide for 2 minutes at 37 °C, washed with PBS supplemented with 100 μg/mL cycloheximide, and then trypsinized with 100 μg/mL cycloheximide at 37 °C. Cells were spun down at 500 x g for 3 minutes at 4 °C, washed twice with cold PBS supplemented with 100 μg/mL cycloheximide. Cells were lysed in polysome lysis buffer (20 mM Tris pH 7.4, 100 mM KCl, 5 mM MgCl2, 1% (v/v) Triton X-100, 100 μg/mL cycloheximide, 500 U/mL RNaseIn Plus, 1x cOmplete protease inhibitor cocktail) and incubated on ice for 10 minutes. The lysate was passed through a 26-guage syringe to ensure efficient cell lysis. The lysate was spun at 1300 x g for 10 minutes at 4 °C and the supernatant was flash frozen with LN2 and stored at −80 °C. The lysate was loaded onto a 10-50% sucrose gradient with cycloheximide and SuperaseIN and centrifuged at 36000x g for 2 hours at 4 °C. Fractionation and absorbance 254 nm measurement was performed using the BioComp gradient fractionator.

---

**Extended Data Figure 1.**
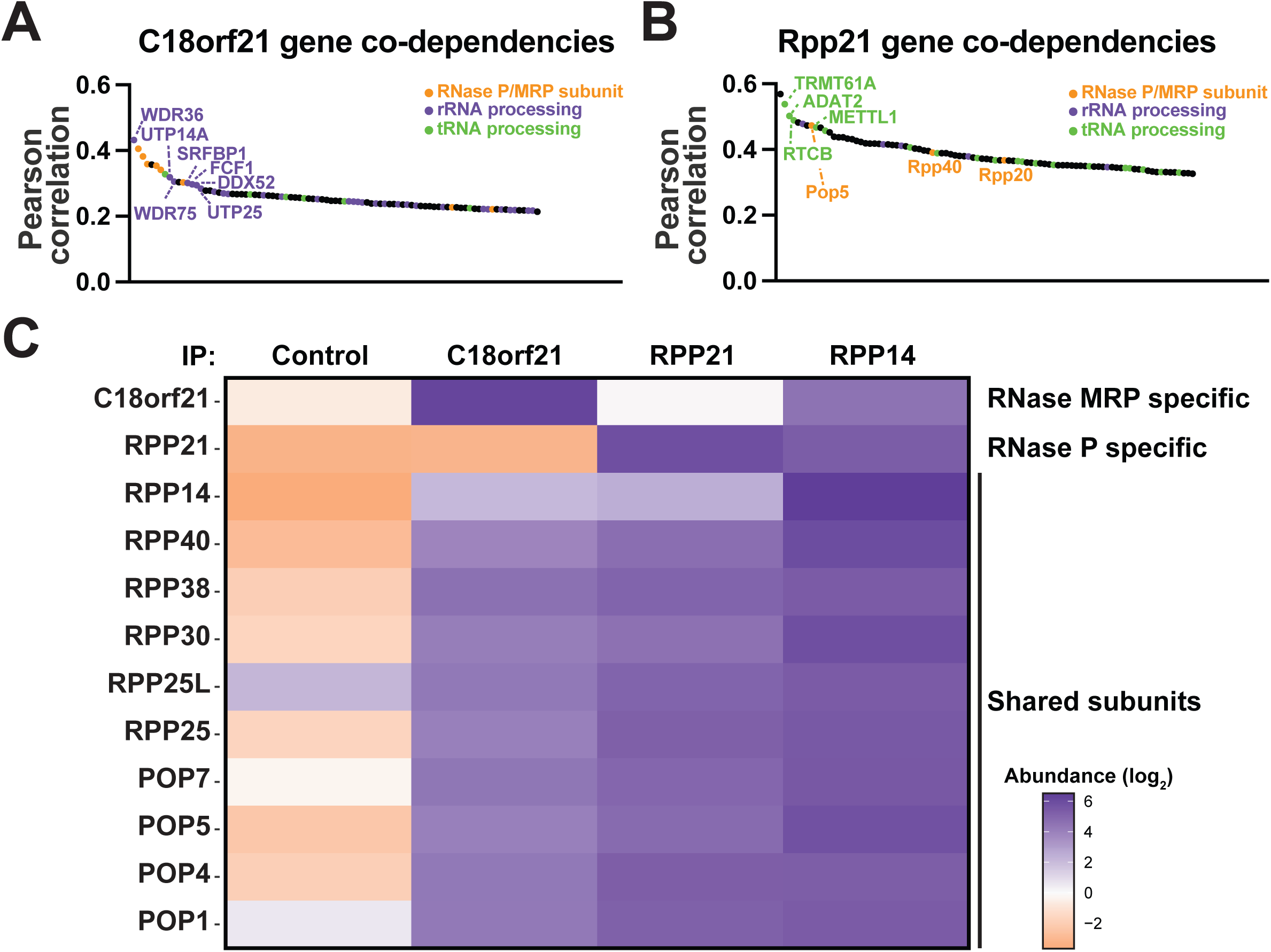
**(A)** A waterfall plot showing the genes with the highest Pearson correlation of CRISPR-Cas9 based targeting effect scores to C18orf21 in the Depmap database. The shared RNase P and MRP components are shown in orange. Genes involved in rRNA processing are shown in cyan and genes involved in tRNA processing are shown in green. **(B)** A waterfall plot showing the genes with the highest Pearson correlation of CRISPR-Cas9 based targeting effect scores to Rpp21 in the Depmap database. The shared RNase P and MRP components are shown in orange. Genes involved in rRNA processing are shown in cyan and genes involved in tRNA processing are shown in green. **(C)** A heat map showing the scaled relative precursor ion abundances of each RNase P/MRP component immunoprecipitated from cell lines ectopically expressing the indicated GFP tagged construct.

**Extended Data Figure 2.**
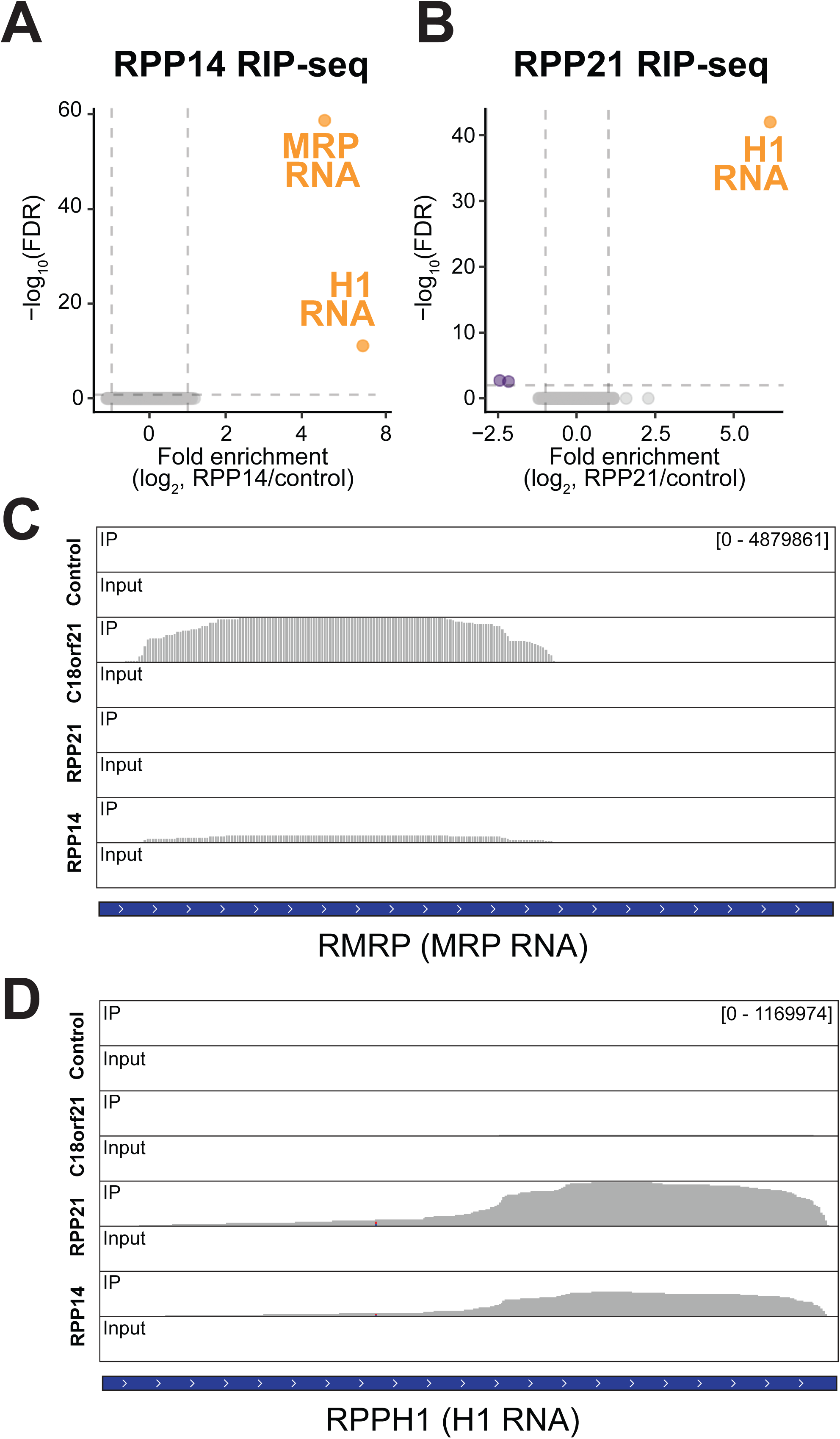
**(A)** A graph showing the Log_10_ False discovery rate (FDR) on the Y-axis and fold enrichment on the X-axis obtained from the indicated ribonucleoprotein-immunoprecipitation (RIP) RNA sequencing experiments performed in duplicate. **(B)** A graph showing the Log_10_ False discovery rate (FDR) on the Y-axis and fold enrichment on the X-axis obtained from ribonucleoprotein-immunoprecipitation (RIP) RNA sequencing experiments performed in duplicate. **(C)** RNA-sequencing coverage plots that display the RMRP reads from the RMRPP1, Rpp14, or Rpp21 immunoprecipitations. The plot for the RMRP reads for RMRPP1 and control cells is duplicated here from figure 3C to allow for comparison between the reads obtained from these immunoprecipitations and the reads obtained from the immunoprecipitations of Rpp14 and Rpp21. **(D)** RNA-sequencing coverage plots that display the RPPH1 reads from the RMRPP1, Rpp14, or Rpp21 immunoprecipitations.

**Extended Data Figure 3.**
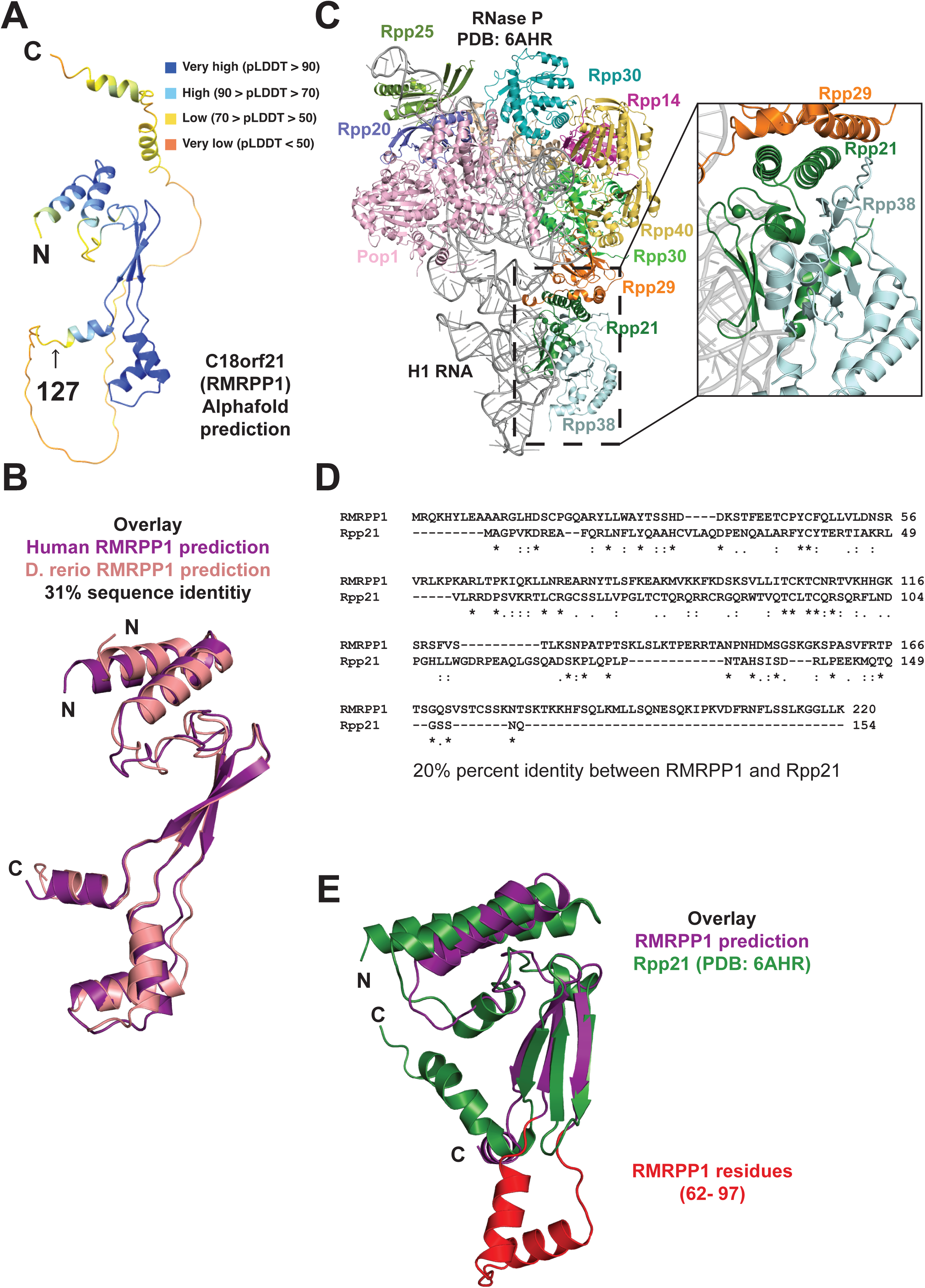
**(A)** The predicted structure of RMRPP1 shown in a cartoon representation. The structure is colored by the predicted local distance difference test (pLDDT) confidence score acquired from the AlphaFold prediction. **(B)** An overlay of the predicted structures of RMRPP1 from human (deep purple) and from *Danio Rerio* (Salmon). **(C)** A cartoon representation of the solved structure of human RNase P PDB ID: 6AHR. Pop5 is shown in wheat, Pop1 is shown in light pink, Rpp40 is shown in yellow orange, Rpp38 is shown in light blue, Rpp30 is shown in teal and green, Rpp29 is shown in orange, Rpp25 is shown in split pea green, Rpp20 is shown in slate blue, Rpp21 is shown in forest green, Rpp14 is shown in magenta, and H1 RNA is shown in grey. The magnified view to the right of the structure highlights the contacts between Rpp21 and Rpp29. **(D)** Sequence alignment of human RMRPP1 and human Rpp21 performed in Clustal omega. **(E)** An overlay of the predicted structure of RMRPP1 (deep purple) and the solved structure of human Rpp21 (Forest green, PDB ID: 6AHR). Residues 62-97 of RMRPP1 are highlighted in red to accentuate the difference in structure between RMRPP1 and Rpp21^45^.

**Extended Data Figure 4.**
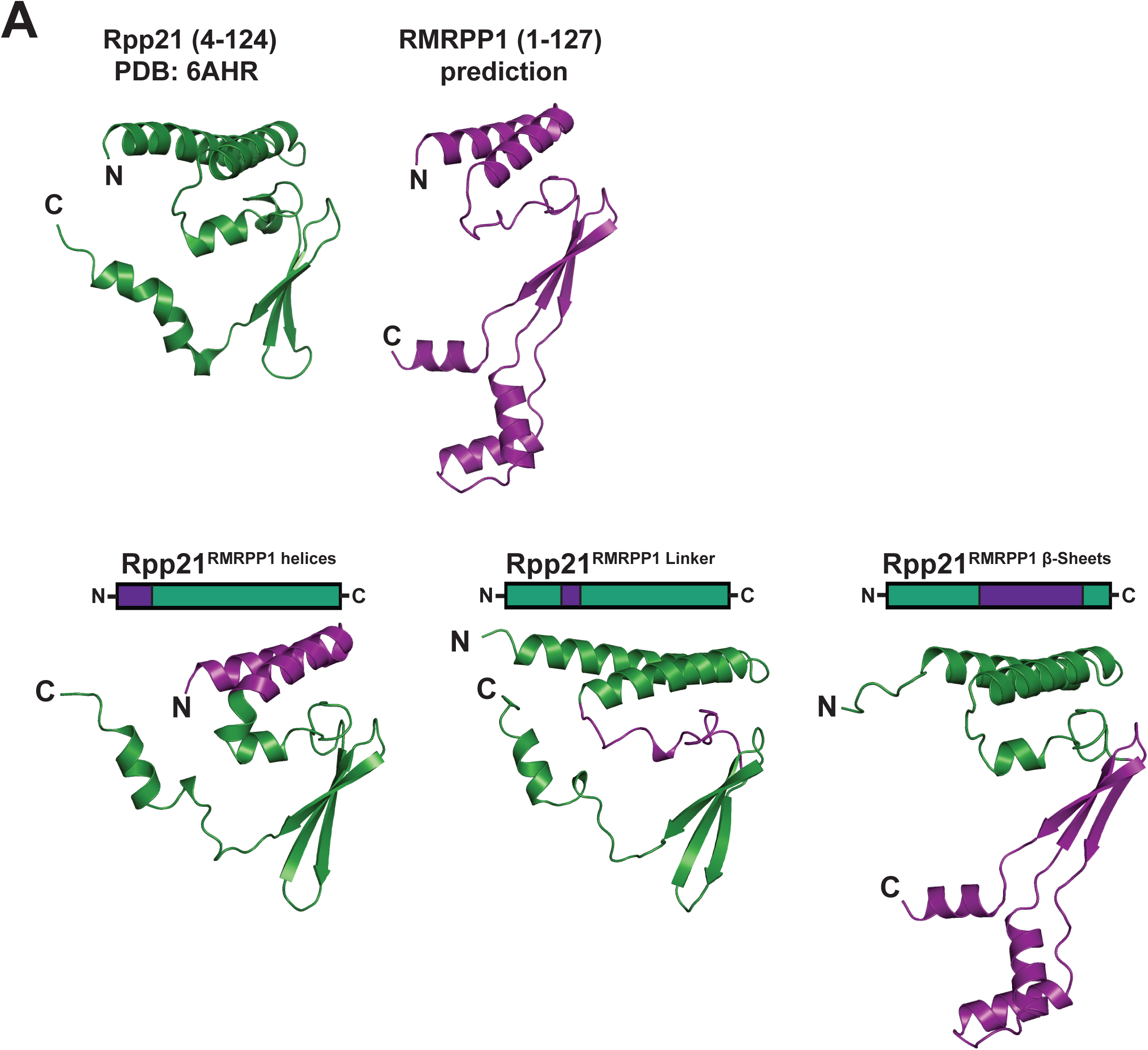
**(A)** Structural predictions for each chimeric construct used in Figure 4. Structural elements taken from RMRPP1 are shown in purple and structural elements taken from Rpp21 are shown in green. Each prediction was done with the full-length construct, for simplicity, only the conserved N-terminal domain is depicted.

**Extended Data Figure 5.**
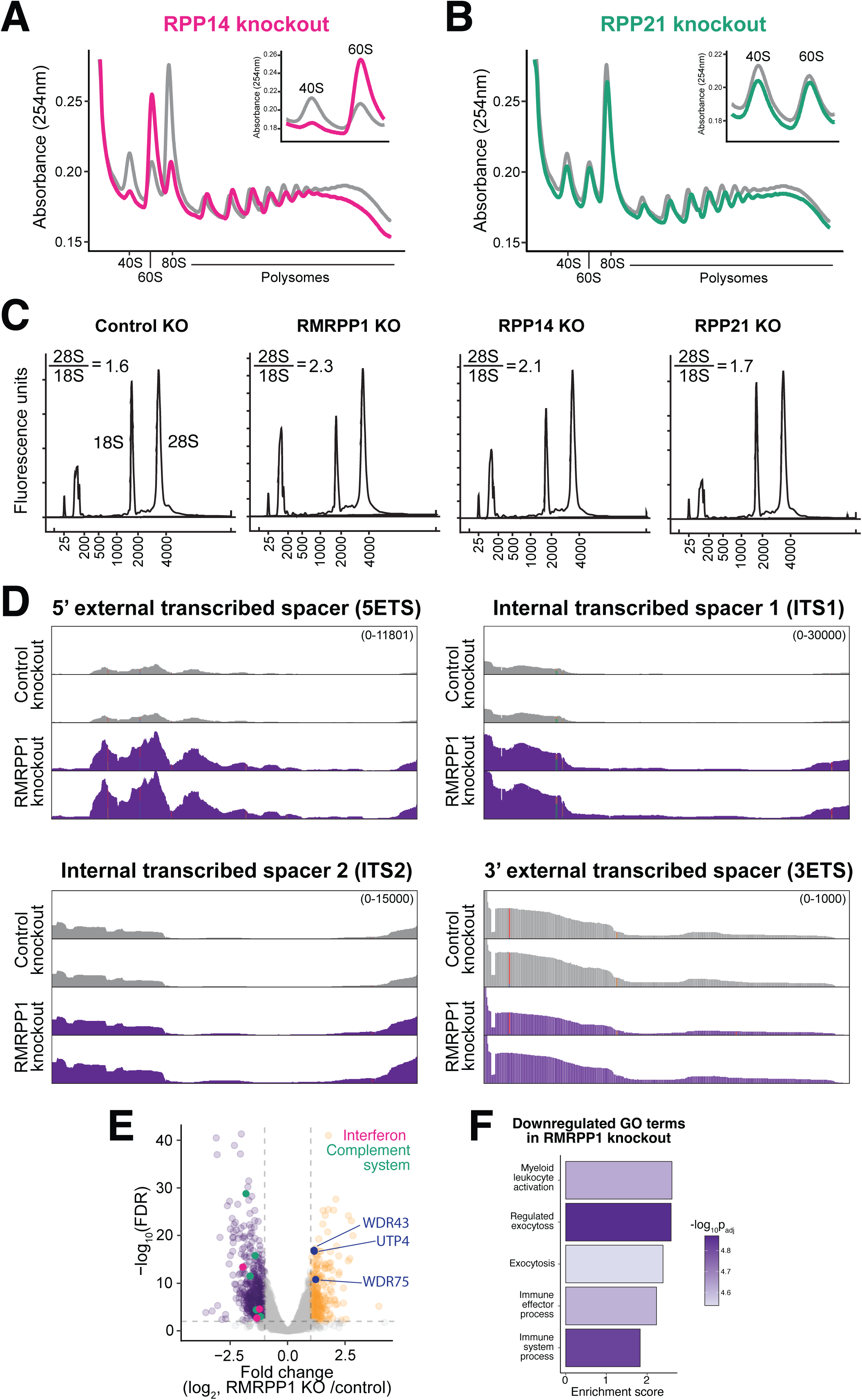
**(A)** Representative polysome profile obtained from the fractionation of cell lysates obtained from Rpp14 knockout cells. The inset in the top right is zoomed in to highlight the 40S to 60S ribosome ratio. **(B)** Representative polysome profile obtained from the fractionation of cell lysates obtained from Rpp21 knockout cells. The inset in the top right is zoomed in to highlight the 40S to 60S ribosome ratio. **(C)** Representative bioanalyzer traces obtained from RNA isolated from control, RMRPP1, Rpp21, or Rpp14 knockout cells. Ratios of 28S:18S were obtained by integrating each peak in the bioanalyzer trace. **(D)** RNA-sequencing coverage plots that display reads for specific regions of the 45S rRNA from the RMRPP1 knockout cells or control knockout cells. (**E**) Volcano plot showing the changes in gene expression in RMRPP1 knockout compared to control knockout cells. Genes with FDR < 0.01 and a fold change > 2 were called as significant. (**F**) Gene set enrichment analysis of RMRPP1 knockout cells to identify gene sets that are downregulated upon RMRPP1 knockout.

**Extended Data Figure 6.**
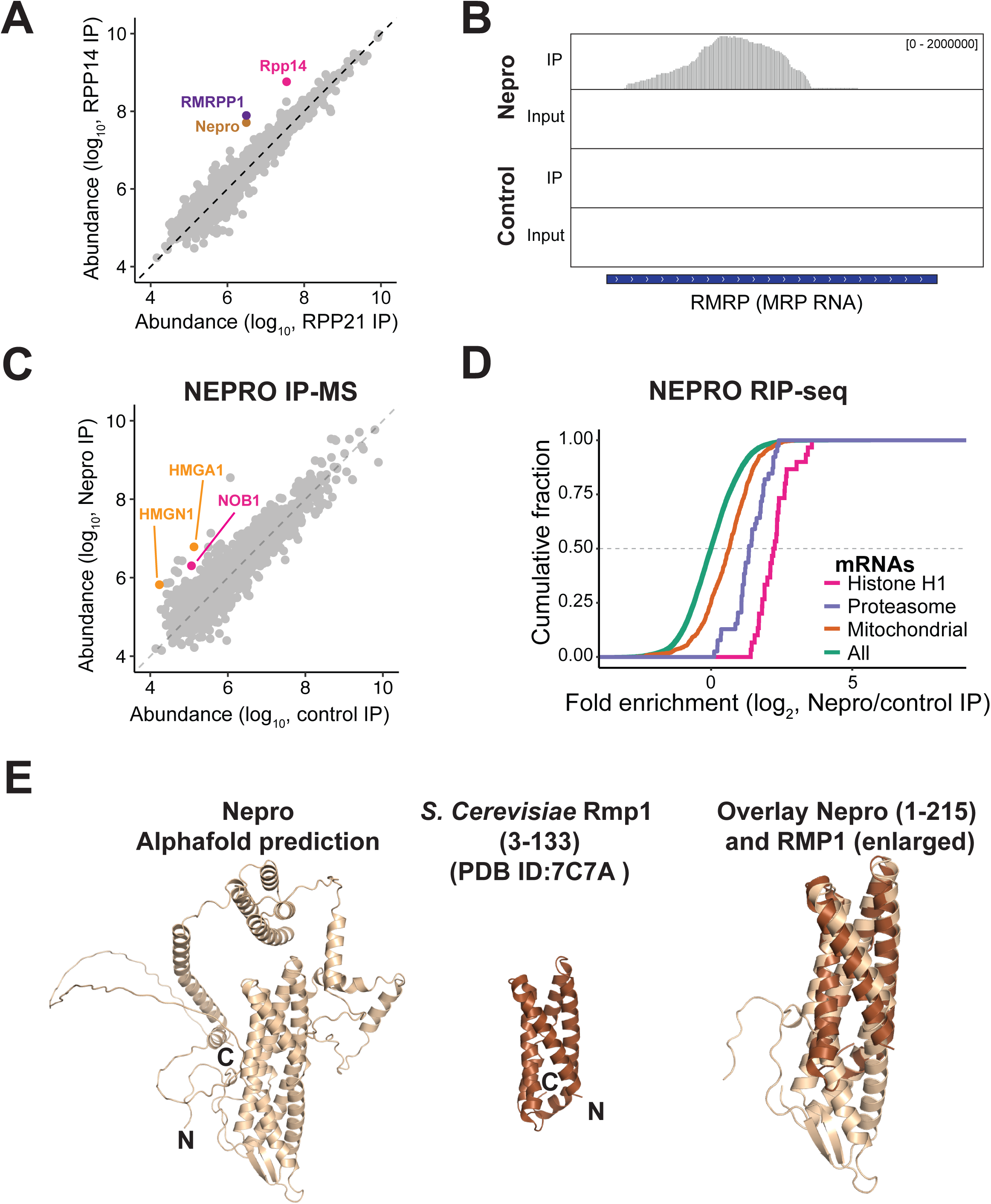
**(A)** A Scatter plot showing the abundance of the proteins detected in the indicated immunoprecipitation-mass spectrometry experiments. Rpp14 is shown in pink, RMRPP1 is highlighted in purple, and Nepro is highlighted in brown. **(B)** RNA-sequencing coverage plots that display reads for RNase MRP RNA. **(C)** The same scatter plot that is shown in Figure 6C highlighting non-RNase MRP/P proteins that were immunoprecipitated by Nepro. **(D)** Plot showing the cumulative fraction on the Y-axis and the log_2_ Fold enrichment on the X-axis of families of mRNA species obtained from ribonucleoprotein-immunoprecipitation (RIP) RNA sequencing experiments. Mitochondrial gene annotations were taken from MitoCarta3.0^50^. **(E)** A cartoon representation of the predicted structure of Nepro that was created using AlphaFold 3 is depicted in wheat and the solved structure of *S. cerevisae* Rmp1 (PDB ID: 7C7A) shown in brown. An overlay of the structurally similar N-terminal helical bundle is shown to the right.

## Notes

### Competing Interest Statement

The authors have declared no competing interest.

